# A rapid microwave method for isolation of genomic DNA and identification of white rot fungi

**DOI:** 10.1101/307066

**Authors:** Ramya G. Rao, A. Ravichandran, A. Dhali, A.P. Kolte, K Giridhar, Sridhar Manpal

## Abstract

White rot fungi (WRF) produce lignolytic enzymes comprised by laccases and peroxidases responsible for mineralization of recalcitrant lignin. Because of the so-called lignin modifying enzymes(LME’s), these fungi have potential applications in biodegradation and bioremediation processes. Increased demand for lignolytic enzymes to exploit their various applications has sparked interest in identifying and characterizing new novel strains of WRF. Despite this undisputed biotechnological significance, molecular identification of WRF, remains a daunting task for researchers as genomic DNA isolation is a tedious process, unsuccessful many a times because of their rigid and resistant cell walls. A rapid, effective and efficient method to identify the innumerable fungal strains within no time is the need of the hour. The fungal mycelia of various unknown as well as know isolates of WRF, after alternative washing with TE buffer and sterile water were suspended in TE buffer. Fungi in solution were then exposed to microwave. The crude extract contained genomic DNA which was extracted and amplified using ITS primers for further identification. Based on sequencing results the identity of known cultures was confirmed, while the unknown cultures were identified as *Clitopilus scyphoides* (*AGUM004*, BankIt2098576 MH172163); Ganoderma *rasinaceum* (*AGUM007*, BankIt2098576 MH172163); *Schizophyllum sp* (*KONA001* BankIt2098576 MH172164; *AGUM011* BankIt2098576 MH172165and *AGUM021* BankIt2098576 MH172166respectively), *Coprinellus disseminatus* (*BANG001*, BankIt2098576 MH172167) and *Lentinus squarrosulus* (*TAMI004*, BankIt2098576 MH172167). The microwave method described for isolating quality DNA of WRF without further purification steps proved a novel method requiring less than ten minutes and minimized the chances of the presence of PCR inhibitors.

**IMPORTANCE:** White rot fungi which decay wood, possess selective lignin degrading enzymes responsible for degrading a wide variety of environmental pollutants, xenobiotic compounds in addition to mineralizing chemicals that are insoluble and recalcitrant. Lignolytic enzymes hold potential towards replacing conventional chemical processes and their increased demand in the market has ignited interest in identifying and characterizing new strains of WRF. A rapid, efficient method capable of quickly identifying fungal isolates is a constraint. The microwave method is a novel quick method for isolating superior quality DNA. Its adoption circumvents the initial purification steps and /or interference of PCR inhibitors, which are encompassed in the use of conventional methods. The microwave method thus permits the thorough amplification of the ITS region thereby aiding in the easy identification of unknown species. Use of the microwave method will permit researchers to obtain DNA from fungi very quickly for further application in molecular studies.

## INTRODUCTION

White rot fungi (WRF) are a group of fungi belonging to the Basidiomycetces which degrade the lignin components from lignocellulosic substances causing bleaching of the wood [1]. WRF produce the set of enzymes *viz.* Laccase (Lac), Manganese peroxidase (MnP), lignin peroxidase (LiP) and Versatile peroxidase (VP) which are responsible for the selective degradation of recalcitrant lignin [2,3]. Because of this lignin modifying enzymes, WRF can degrade wide varieties of environmental pollutants, xenobiotic compounds and also mineralize chemicals that are insoluble and recalcitrant [4]. Hence, they have potential applications in biodegradation and bioremediation processes. The ability of WRF to degrade lignocellulosic, a central aspect in industrial uses of cellulosic biomass, such as bioethanol production, manufacture of cellulose based chemicals and materials including paper and recently in crop residues as animal feed to improve its nutritive value as they promise environmental friendly technologies [5,6]. Their biotechnological significance has caused a drastic increase in the demand of these enzymes in the recent few decades.

Limiting amounts of lignolytic enzymes however, are produced by WRF, and identification of the produced enzymes impedes their commercial use in innumerable potential applications. The species level identification of WRF (microorganisms), provides deeper insights on fungal life cycle, evaluation and molecular aspects of the protein production which in turn helps researchers to enhance the production of enzymes, identification of new species and meet the increased demand [7]. On the other hand, we lack standardized protocols for conducting routine molecular biology research of these microorganisms. Due to high polysaccharide contents, the cell walls of WRF are rigid and are resistant to DNA extraction by traditional methods [8]. In addition, methods involved in DNA isolation are laborious, tricky, time consuming and very expensive for isolating DNA of excellent quality [9]. All these methods commonly employ the use of detergents such as SDS for cell wall lysis, which often inhibits further purification manipulations [10]. Most of these methods involve innumerable steps that take lot of time and in addition possess the threat of contributing PCR inhibitors.

In the present study, we report a simple and rapid method based on the application of microwave for DNA isolation from some of the wild isolates of WRF which was then used for PCR amplification and species identification of the unknown strains of WRF. This method has also been compared with other easy and rapid methods being used for different fungal species by researchers around the world. Qualitative and quantitative analysis of DNA extracted from different methods was evaluated based on yield of DNA, purity in terms of A260/A280 ratio, PCR and gel electrophoresis. Unknown WRF isolated were identified by sequencing the PCR product of genomic DNA obtained from microwave method.

## MATERIALS AND METHODS

### Isolation and storage of WRF strains

Fruiting bodies or basidiocarps of WRF were collected in clean dry self-sealing polythene bags from forest areas. Amongst the seven wild fungal isolates, KONA001was collected from 13.8048 º N, 75.2530 º E; AGUM004, AGUM007, AGUM011 and AGUM021 were collected from 13.5187 º N, 75.0905 º E; BANG001 was collected from12.9470 º N, 77.6077 º E while TAMI004 was collected from location 08.9342 º N, 77.2778 º E from Karnataka, India. In all cases the substrate was represented by wood found in various stages of decay. Pure cultures from collected samples were obtained by tissue culture technique [11]. All pure cultures were maintained on PDA slants and stored at 4°C until further use. The cultures were marked with information such as number and procurement location. *Coriolus versicolor* (MTCC138), *Ganoderma lucidium* (MTCC1039) and *Pleurotus sajorcaju* (MTCC141) obtained from Microbial Typing Culture Collection, Chandigarh, India were used as the reference cultures.

### DNA extraction

Four different methods were evaluated for the Extraction of DNA from the selected unknown wild isolates of WRF:

### Method 1: Rapid mini preparation of DNA

The rapid mini preparation of DNA [12] method was comprised of a small amount of revived culture being suspended in 500μL of lysis buffer containing 400mM Tris-HCl (pH8), 60mM EDTA(pH8), 1% Sodium dodecyl sulphate(SDS), 150mM NaCl and the lumps disrupted using sterile loop. Samples were incubated for 10 minutes at room temperature. The samples were mixed with potassium acetate (pH 4.8) and centrifuged at10000Xg for 2 min and the supernatant in fresh Eppendorf spun again. Then the supernatant was mixed with an equal volume of isopropyl alcohol by brief inversion. The sample was centrifuged for 2 minutes at 10,000Xg and the supernatant was discarded. The pellet had DNA and was washed in 300μL 70% alcohol. After the pellet was centrifuged for 1 minute at, the supernatant was discarded, and DNA was air dried. The isolated DNA was then dissolved in 50μl 1X TE buffer.1μL DNA suspension was used for PCR.

### Method 2: Thermolysis Method

In the thermolysis method [13] a small quantity of mycelia was picked by help of a sterile needle from the fully-grown culture and transferred into 100μL sterile water in a 2 mL micro centrifuge tubes. The mixture was thoroughly vortexed and centrifuged at 10,000g for 1 minute.to the pellet, after discarding supernatant 100μl lysis was added. The mixture was incubated at 85 C in a water bath for 25 minutes. The crude extract contained genomic DNA.1μl supernatant was used for PCR.

### Method 3: Microwave thermal shock method

As per the microwave thermal shock method [14] a small quantity of each of the revived cultures was suspended in 1mL of washing solution containing 50mmol L^-1^ Tris-HCl, pH 7.7, 25mmol L^-1^ EDTA, 0.1% Sodium dodecyl sulphate (SDS), 0.1% polyvinylpyrrolidone (PVP). Samples were centrifuged at 6000Xg for 1min. and the pellets were resuspended in 35μL of lysis buffer containing 50mmol L^-1^ Tris-HCl, pH 8, 25mmol L^-1^ EDTA, 3% SDS, 1.2% PVP. The mixture was then placed in a microwave oven (Electrolux EK30CBB6-MGZ; RF output-900W) and heated at 700W for 45s. 400μl of pre-warmed extraction solution containing 10mmol L^-1^ Tris -HCl, pH 8, 1mmol L^-1^ EDTA, 0.3 mol L^-1^ Sodium acetate, 1.2% PVP were added to the microwaved sample. The DNA was extracted using phenol chloroform solution followed by isopropyl alcohol precipitation and 70% ethanol wash. Precipitated DNA was then resuspended in 100μL TE buffer (pH 8.0). One μL buffer was used for PCR.

### Method 4: Microwave method

All the selected wild isolates were removed from storage and revived on PDA slants at 27±2°C for 7-10 days. A small amount of mycelium from the grown culture was picked with the help of a sterile needle and transferred into 1000μL of 1XTE in 2mL micro centrifuge tubes. The mixture was thoroughly vortexed and centrifuged at 10,000Xg for 1min. The supernatant was discarded, and the pellet was washed with 1000μL of 1XTE again followed by with 1000μl of sterile water. After the wash, pellet obtained was resuspended in 200μl of 1XTE. The mixture was then placed in a microwave oven (Electrolux EK30CBB6-MGZ; RF output-900W) and heated at 900W for 1min. twice. The crude extract contained genomic DNA. 1μl supernatant was used for PCR.

The quality and quantity of all isolated DNA was checked using a Nano drop (Thermo Scientific) in terms of A260/A280 ratio and ηg /μL respectively [15].

### Amplification of ITS regions of DNA

Each PCR mixture contained, 10μL Master Mix (Thermo scientific), 0.5 μL of forward and reverse primers each, and 8μL of nuclease free water and 1μL of DNA template to be amplified. The primer base pairs used for the amplification of ITS regions were: forward primer ITS1F (CTTGGTCATTTAGAGGAAGTAA) and reverse primer ITS4 (TCCTCCGCTTA TTG ATA TGC) [16]. Primers were procured from Eurofins, India. The PCR consisted of an initial denaturing step of 5min at 95 ºC followed by 35 cycles of 94 ºC for 50s, 54 ºC for 50s and 72 ºC for 50s and finished by final extension step for 10 minutes at 72 ºC [17]. Amplified PCR products were resolved by electrophoresis through 1% agarose gel stained with ethidium bromide.

The PCR products of the seven unknown cultures KONA001, AGUM004, AGUM007, AGUM011, AGUM021, BANG001, TAMI004 and MTCC culture MTCC138 were given for sequencing. Sequences obtained from Eurofins India were aligned against EMBL DNA database. All sequences were then checked against Gene bank with the help of BLAST. Culture names were assigned based on more than 99% sequence similarity [18].

### Statistical Analysis

ANOVA was performed to compare the different DNA isolation methods within each WRF isolate for DNA yield as well as purity. Mean and standard deviations were determined for replicates. For all the statistical analysis, software, SAS 9.3 was used.

## RESULTS

The four different methods were used to extract DNA from 10 different WRF (KONA001, AGUM004, AGUM007, AGUM011, AGUM021, BANG001, TAMI004, MTCC138, MTCC1039, and MTCC149). The yields of DNA and quality of DNA, in terms of A260/A280 ratio, obtained from different methods are significantly different at confidence interval 99 % (Table1). Concentration of DNA in case of method 4 is less than that of three methods 1, 2 and 3. However quality is superior in case of DNA isolated from Method 4 as compared to the other three methods (Table 1). Concentrations of DNA and purity of DNA obtained by all the methods are in acceptable range for further molecular studies.

**Table 1.** Concentration (yield (ηg/μl)) and Purity of DNA isolated from different WRF using four methods

| WRF | DNA Yield ($\eta\text{g}/\mu\text{l}$) | | | | $A_{260}/A_{280}$ Ratio | | | |
| --- | --- | --- | --- | --- | --- | --- | --- | --- |
|  | Method1 | Method2 | Method3 | Method4 | Method1 | Method2 | Method3 | Method4 |
| MTCC138 | 66.9±1.29 | 61.9±1.49 | 64.9±0.58 | 58.8±0.69 | 1.93±0.02 | 2.01±0.01 | 1.93±0.03 | 2.03±0.04 |
| MTCC149 | 110.7±1.52 | 110.5±0.7 | 114.5±1.8 | 107±1.45 | 2.09±0.07 | 2.11±0.01 | 2.11±0.04 | 2.1±0.02 |
| MTCC1039 | 87.5±1.22 | 84.3±1.37 | 85±1.43 | 79.6±0.78 | 2.04±0.06 | 2.05±0.02 | 1.99±0.02 | 2.03±0.05 |
| AGUM004 | 238.1±1.64 | 226.3±3.71 | 229.5±1.62 | 227.5±0.93 | 2.07±0.04 | 2.1±0.02 | 2.1±0.01 | 2.06±0.07 |
| AGUM007 | 128.1±1.78 | 120.9±1.2 | 127.3±1.43 | 126.5±1.58 | 2±0.03 | 2.04±0.03 | 2.03±0.01 | 2.04±0.04 |
| AGUM011 | 339.7±0.91 | 334.8±2.52 | 341.2±1.63 | 338.6±0.77 | 1.88±0.02 | 1.88±0.04 | 1.85±0.02 | 1.88±0.04 |
| AGUM021 | 122.3±1.87 | 126.5±2.18 | 136.2±1.34 | 119±1.43 | 2.04±0.04 | 2.03±0.03 | 2.04±0.03 | 2.06±0.06 |
| KONA001 | 152.8±2.62 | 144.9±2.36 | 159.7±1.23 | 137±1.97 | 1.92±0.02 | 2.02±0.02 | 1.95±0.02 | 2.02±0.02 |
| BANG001 | 165.3±1.07 | 147±2.31 | 157.2±0.94 | 142±1.46 | 2±0.02 | 2.09±0.03 | 2.05±0.05 | 2.05±0.04 |
| TAMI004 | 99.8±0.3 | 94.4±1.35 | 99.7±2.77 | 93±0.95 | 2.02±0.03 | 2.07±0.01 | 2.03±0.01 | 2.05±0.04 |

The PCR amplification and gel electrophoresis (Fig. 1) reveals that using method 1and 2, only one sample each was got amplified and method 3, only two samples were amplified (Fig. 1 A, B and C respectively). Only DNA isolated for all the 10 samples from method 4 was subjected to amplification (Fig1 D). PCR products obtained with the help of method 4 for the seven unknown cultures and one MTCC culture *Coriolus versicolor* (MTCC138) were sequenced and were identified. The unknown cultures submitted to GenBank were identified as *Clitopilus scyphoides* (*AGUM004*, Bank It 2098576 MH172163); Ganoderma *rasinaceum* (*AGUM007*, BankIt2098576 MH172163); *Schizophyllum sp* (*KONA001* Bank It 2098576 MH172164; *AGUM011* BankIt 2098576 MH172165and *AGUM021* BankIt2098576 MH172166respectively), *Coprinellus disseminatus* (*BANG001*, BankIt2098576 MH172167) and *Lentinus squarrosulus* (*TAMI004*, BankIt2098576 MH172167).

**Fig 1.**
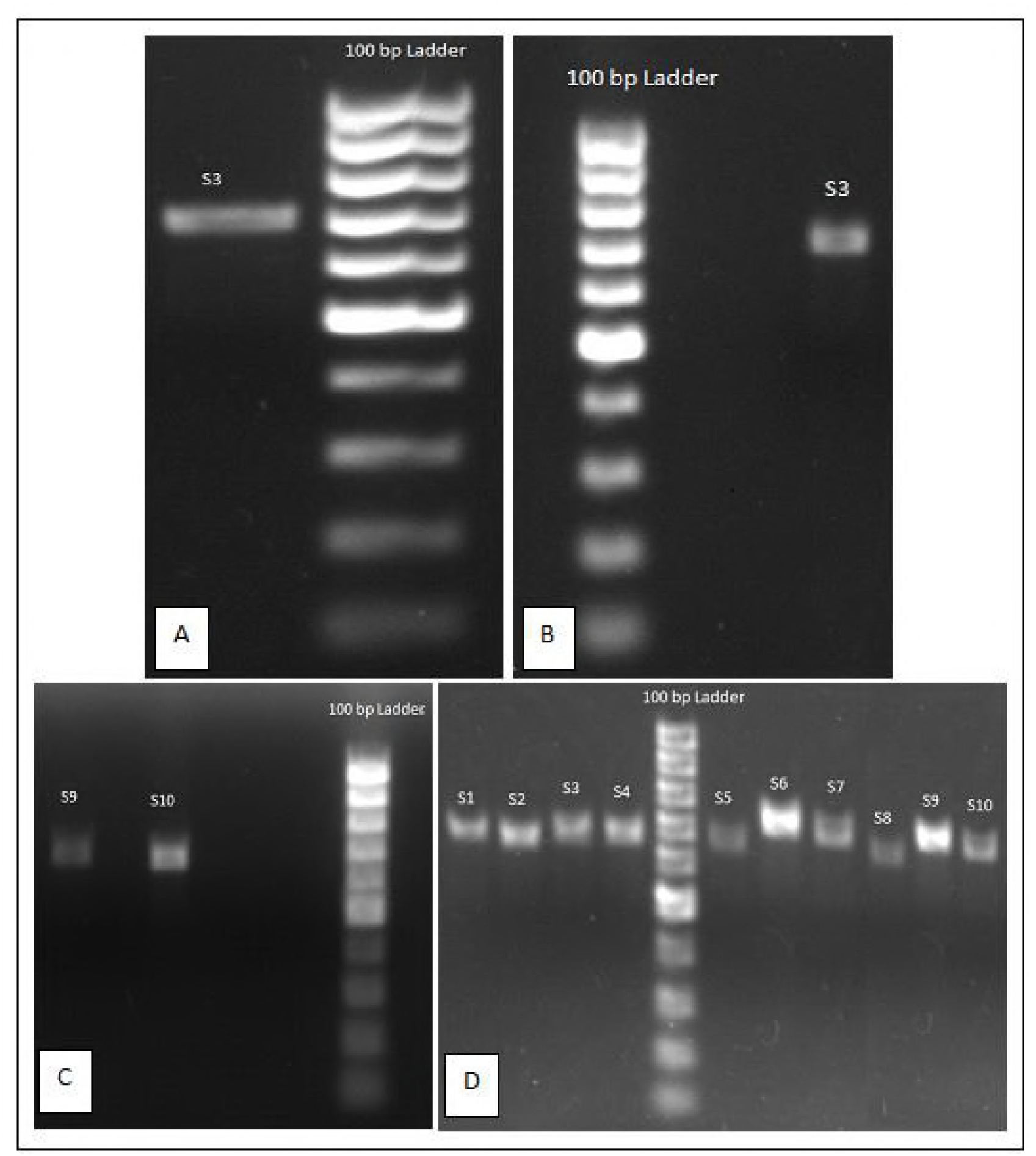
Agarose gel electrophoresis images for PCR amplifications of ITS region of genomic DNA of different WRF isolated using Method 1(A), Method 2(B), Method 3 (C) and Method 4 (D)Samples S1-10 are from WRF KONA001, AGUM004, AGUM007, AGUM011, AGUM021, BANG001, TAMI004, MTCC138, MTCC149 and MTCC1039.Lanes for samples which were not amplified are not shown in the gel images.

## DISCUSSION

Though all the four methods used for isolating genomic DNA are capable of yielding DNA of acceptable quality and quantity, only DNA obtained by help of the microwave method could get amplified in PCR. The major reason behind the DNA not being amplified was the presence of PCR inhibitors [19]. PCR may be inhibited by the presence of certain chemicals / biomolecules released from fungal species which may vary from species to species, growth status and media used for cultivation [20].

The microwave method offers several advantages. As this method takes less than 10 minutes to isolate DNA bulk identification of WRF strains can be achieved very quickly saving precious time by avoiding innumerable cumbersome steps as is in case of the other methods. Indeed, this is the first report to isolate WRF genomic DNA by microwave method. In other protocols, a microwave method was reported for bacterial genomic DNA [21, 22]. The DNA isolated does not require chemicals like phenol or chloroform. This method also prevents the release of cell wall chemicals of WRF and other chemicals released from the species which are known to be potent PCR inhibitors. The yield of DNA and its purity is also in acceptable range and proven to amplify ITS region and intern’s species level identification.

The microwave method described here for WRF is a novel method that takes less than 10 minutes to isolate DNA without any initial purification steps and /or interference of PCR inhibitors, permitting the amplification of the ITS region and thereby enabling the easy identification of unknown species.

Future works need to be carried out in the direction of other molecular biology research with the isolated DNA such as whether screening of genes of interest, cloning and expression in a different host for increased yield of proteins/enzymes which are of commercial and clinical importance, phylogenetic tree construction etc., are possible. There is a possibility of using Microwave method in environmental and biotechnological studies, because rapid DNA isolation gives a simple solution to sequence several strains directly or by micro arrays [22].

## CONCLUSION

The microwave method is a novel method taking less than ten minutes to isolate superior quality DNA. Its adoption circumvents the initial purification steps and /or interference of PCR inhibitors, which are encompassed in the use of conventional methods. The microwave method thus permits the thorough amplification of the ITS region thereby aiding in the easy identification of unknown species. Further work in the direction of supplemental molecular biology research with the isolated DNA such as screening for genes of interest, cloning and expression in a different host for increased yield of proteins/enzymes of commercial and clinical importance, phylogenetic tree construction etc., are however warranted.

## ACKNOWLEDGEMENT

The financial assistance (GrantNo.BT/PR11205/AAQ/1/589/2014) provided by Department of Biotechnology, (DBT), Government of India, New Delhi, is gratefully acknowledged by the authors. The authors thank the Director, National Institute of Animal Nutrition & Physiology, for providing all the facilities for conduct of the research work.

## Author contribution statement

RRG executed this work. RA contributed to the design, AD and APK assisted in execution, and analysis of this work. MS and GK contributed to the interpretation of results and drafting the submission.

